# *ANTHRACNOSE RESISTANCE GENE2* confers fungal resistance in sorghum

**DOI:** 10.1101/2022.06.24.497546

**Authors:** Demeke B. Mewa, Sanghun Lee, Chao-Jan Liao, Augusto M. Souza, Adedayo Adeyanju, Matthew Helm, Damon Lisch, Tesfaye Mengiste

**Author notes:** Corresponding author: Tesfaye Mengiste.

## Abstract

Sorghum is an important food and feed crop globally; its production is hampered by anthracnose disease caused by the fungal pathogen *Colletotrichum sublineola* (*Cs*). Here, we report identification and characterization of *ANTHRACNOSE RESISTANCE GENE 2* (*ARG2*) encoding a nucleotide-binding leucine-rich repeat (NLR) protein that confers race-specific resistance to *Cs* strains. *ARG2* is one of a cluster of several *NLR* genes in the sorghum differential line SC328C that is resistant to some *Cs* strains. This cluster shows structural and copy number variations in different sorghum genotypes. Different sorghum variants carrying independent *ARG2* alleles provided the genetic validation for the identity of the *ARG2* gene. *ARG2* expression is induced by *Cs*, and chitin induce *ARG2* expression in resistant but not in susceptible lines. ARG2-mediated resistance is accompanied by higher expression of defense and secondary metabolite genes at early stages of infection, and anthocyanin and zeatin metabolisms are upregulated in resistant plants. Interestingly, ARG2 localizes to the plasma membrane when transiently expressed in *Nicotiana benthamiana*. Importantly, *ARG2* plants produced higher shoot dry matter than near-isogenic lines carrying the susceptible allele suggesting absence of an *ARG2* associated growth trade-off. Further, ARG2-mediated resistance is stable at a wide range of temperatures. Our observations open avenues for resistance breeding and for dissecting mechanisms of resistance.

## Introduction

Sorghum [*Sorghum bicolor* (*L.*) *Moench*] is a crop of global importance that used for food, feed and biofuel. The productivity of sorghum is constrained by anthracnose diseases caused by the fungus *Colletotrichum sublineola* (*Cs*), which results in significant crop losses (Thakur, 2007; Little and Perumal, 2019). Although sorghum disease resistant germplasm is widely available, the molecular genetics of anthracnose resistance has been poorly studied. Generally, plant immune pathways are placed into two categories based primarily on pathogen derived molecules that are perceived and the kinetics of immune responses (Dangl et al., 2000). Recognition of pathogen derived signature molecules, pathogen associated molecular patterns (PAMPs), at the cell surface is mediated by pattern recognition receptors triggering Pattern Triggered Immunity (PTI). PTI is associated with varying degrees of quantitative resistance (Godfrey and Rathjen, 2001). Accumulation of phytoalexin 3-deoxanthocyanidin at the site of infection has been linked to quantitative resistance to *Cs* (Tenkouano et al., 1993; Lo et al., 1999; Cui et al., 2015). The second pathway is mediated through the intracellular recognition of pathogen effectors by nucleotide-binding leucine-rich repeat (NLRs) proteins, which initiate effector trigger immunity (ETI), an effective and often highly specific disease resistance (Rose et al., 2004). ETI is heightened immunity accompanied by the hypersensitive response (HR) (Boyes et al., 1998). The highly polymorphic nature of NLRs and the strong selective pressure on pathogens drives an arms race between plants and pathogens. Resistance to anthracnose involves both qualitative and quantitative resistance mechanisms (Patil et al., 2017; Nelson et al., 2018).

*NLR* genes are widely used in resistance breeding of crops and have been studied in different systems (Wang et al., 2017; Lagudah and Periyannan, 2018). The high penetrance of NLR regulated resistance makes resistance breeding easier but it is also prone to overcome by virulent pathogens. The sorghum genome encodes hundreds of putative *NLR*s an estimated 97% of which occur in clusters (Cheng et al., 2010). In contrast, quantitative resistance is known to be durable and broad-spectrum but confers only partial resistance, resulting in a better yield in infected fields rather than complete prevention of disease (St. Clair, 2010). Sorghum variants with race specific or quantitative anthracnose resistance have been reported (Cuevas et al., 2014; Burrell et al., 2015; Felderhoff et al., 2016; Patil et al., 2017). However, studies focused on the overall genetic architecture of resistance rather than the identity of individual genes (Nelson et al., 2018).

The interaction between the rice blast fungus *Magnaporthe oryzae,* and rice (*Oryza sativa*) appears to closely mirror sorghum interaction with *Cs*. In both cases, the infection starts with a biotrophic phase followed by a necrotrophic phase. The genetic control and molecular mechanisms of resistance to rice blast disease have been well studied. More than 30 rice blast resistance genes and 12 *M. oryzae* effectors have been identified (Wang et al., 2017). Most of these rice genes encode NLRs that confer race-specific resistance. There have been few studies on molecular mechanisms and specific genes that regulate sorghum resistance to this pathogen. Recently, the *ARG1* gene encoding a sorghum NLR was shown to confer broad-spectrum resistance to anthracnose (Lee et al., 2022). Global mRNA and microRNA expression dynamics in response to anthracnose define components of sorghum responses to the pathogen (Fu et al., 2020).

This study was initiated to identify sorghum anthracnose resistance genes and determine their functions in resistance. We report the identification of the *ARG2* gene, encoding a canonical NLR protein that confers complete resistance to some *Cs* strains. The identification of *ARG2* is supported by multiple variants that carry resistant or susceptible *ARG2* alleles that correspond to their resistance or susceptible responses to *Cs* strains. The genomic and molecular characterization of *ARG2* locus, and impact of *ARG2* on defense gene expression, accumulation of secondary metabolites, pathogen resistance at elevated temperatures, and defense growth tradeoff is determined.

## Results

### Identification of resistant sorghum lines

Twenty-five sorghum lines were screened using multiple strains of *Colletotrichum sublineola* (*Cs*) to identify resistant genotypes (Supplemental Table S1). SC328C displayed a clear-cut resistance to *Cs* strains Csgl1 and Csgrg and susceptibility to the strains Csgl2, Cs27 and Cs29. SC328C is part of a collection of sorghum differential lines that have been used for race identification, and its resistance to other strains of *Cs* were reported in previous studies (Moore et al., 2008; Prom et al., 2012). Due to the strong resistance phenotype in SC328C, the genetic control of this resistance was studied in detail.

In detached leaf assays, TAM428 displayed enhanced disease symptoms with expanding and coalescing chlorotic and necrotic lesions starting at the site of inoculation with Csgl1 (Figure 1A). The pathogen proliferated and severely damaged TAM428 leaf tissue. The leaf tissue of SC328C at the site of drop inoculation with Csgl1 became brown but the tissue remained intact and there were no disease symptoms (Figure 1A). In susceptible genotypes, disease symptoms generally appeared in five to seven days after spray inoculation. Pathogen growth was observed as the production of acervuli (fungal reproductive structures), with setae that appear as black spots marking susceptibility (Figure 1B). Staining of the inoculated tissue with WGA-AF488, which stains the fungal hyphae (Redkar et al., 2018), revealed proliferous acervuli on TAM428 but not on SC328C (Figure 1B). Similarly, trypan blue staining of infected leaf tissue revealed a dense mass of mycelia on TAM428 whereas no pathogen growth occurred on SC328C (Supplemental Figure S1A). Thus, inoculated leaf tissue showed enhanced fungal growth on susceptible lines but no fungal growth on resistant lines. After spray-inoculation, the susceptible genotype TAM428 showed widespread disease lesions and enhanced pathogen growth on multiple leaves per plant (Figure 1C). The disease lesions in TAM428 often cover more than half the area of growing leaves and the entire leaf sheath. At 7-10 days post inoculation, plant responses ranged from widespread necrotic lesions to the complete collapse of infected leaves or the entire plant whereas SC328C showed only HR-like responses. The sorghum reference line BTx623 was also susceptible to Csgl1 while in the sweet sorghum reference line Rio resistance was similar to that of SC328C (Figure 1C). Pronounced anthracnose symptoms appear on leaf sheath of susceptible plants when the whole plant was spray-inoculated, and the leaf tissue often tears apart along the leaf venation (Figure 1D).

**Figure 1.**
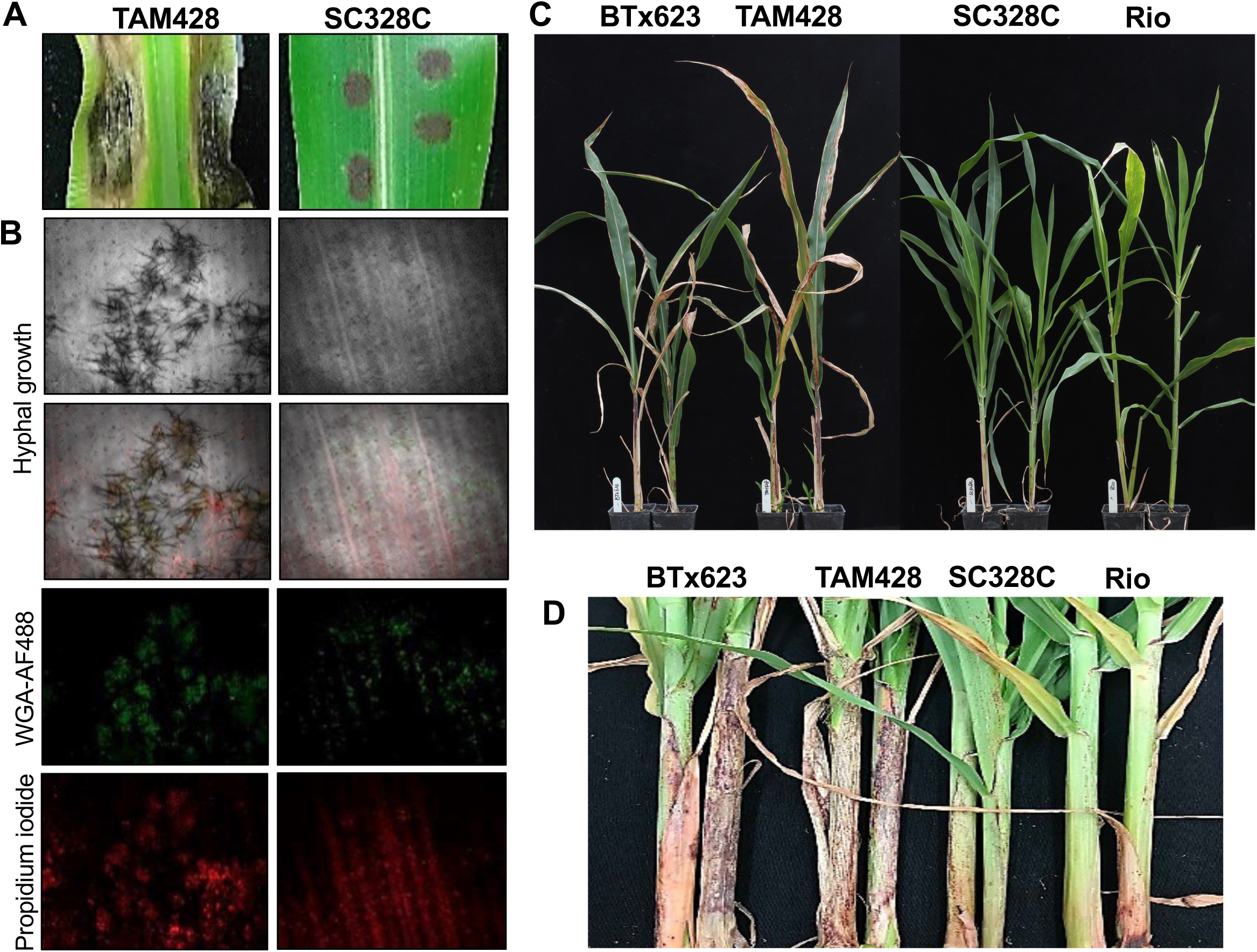
Disease symptoms and fungal growth in sorghum variants. **(A)** Disease symptom and resistance responses at 7 days after drop-inoculation (dai) of *Colletotrichum sublineola* (*Cs*) strains Csgl1 conidial suspension (20 µL, 1x10^6^ conidia/mL) on detached leaves. **(B)** Fungal growth in drop inoculated leaves. Inoculated tissues with *Cs* Csgl1 were stained WGA-AF488 and propidium iodide at 6 dai and examined using a confocal microscope. WGA-AF488 (green) stains fungal hyphae, and propidium iodide (red) stains dead host cells. Pathogen acervuli are visible on TAM428 but not on SC328C which show no pathogen growth. **(C)** Disease symptoms on whole plants at 14 days after spray-inoculation. The conidial suspension (*Cs* Csgl1; 1x10^6^ conidia/mL) was uniformly spray-inoculated on sorghum plants. **(D)** Disease symptom and resistance response on leaf sheath at 15 days after spray-inoculation of the whole plant.

The disease resistance in SC328C was also effective under high temperature (38°C) in a detached-leaf disease assay (Supplemental Figure S1B). The resistance to Csgl1 in SC328C was not compromised at 38°C whereas the symptoms in the susceptible line BTx623 was delayed by several days. Similar observations were made with resistant or susceptible near-isogenic lines (NILs) (Supplemental Figure S1C). Detached leaves of TAM428 did not survive at 38°C.

### A single dominant locus in SC328C regulates resistance

To determine the inheritance of anthracnose resistance in SC328C to *Cs*, SC328C was crossed to two susceptible lines, TAM428 and BTx623 (Supplemental Table S1). The F_1_ progenies of both crosses were resistant, as were the F_1_ progenies of the reciprocal SC328C x TAM428 cross (Figure 2A). The F_2_ progenies derived from self-fertilization of the BTx623 x SC328C F_1_ plants resulted in 298 resistant and 91 susceptible progenies, which fitted to a 3:1 (X^2^=0.464, *P*>0.05) phenotypic ratio. Similarly, evaluation of 1,131 F_2_ progenies of the TAM428 x SC328C cross resulted in 853 resistant and 278 susceptible plants that also fitted well to a 3:1 (X^2^=0.704, *P*>0.05) phenotypic ratio. The disease symptoms on BTx623 appeared later and were less severe compared to that of TAM428 (Figure 1C). Therefore, we used the TAM428 x SC328C population for gene identification because of the enhanced penetrance of the disease phenotypes in this genetic background. The parental lines and progenies showed consistent disease responses to the two *Cs* strains Csgl1 and Csgrg suggesting that the resistance to these two *Cs* fungal strains is regulated by the same locus. All the above genetic analyses strongly suggested that a single dominant locus in SC328C confers resistance to both strains, hereafter designated as *ANTHRACNOSE RESISTANCE GENE2* (*ARG2*).

**Figure 2.**
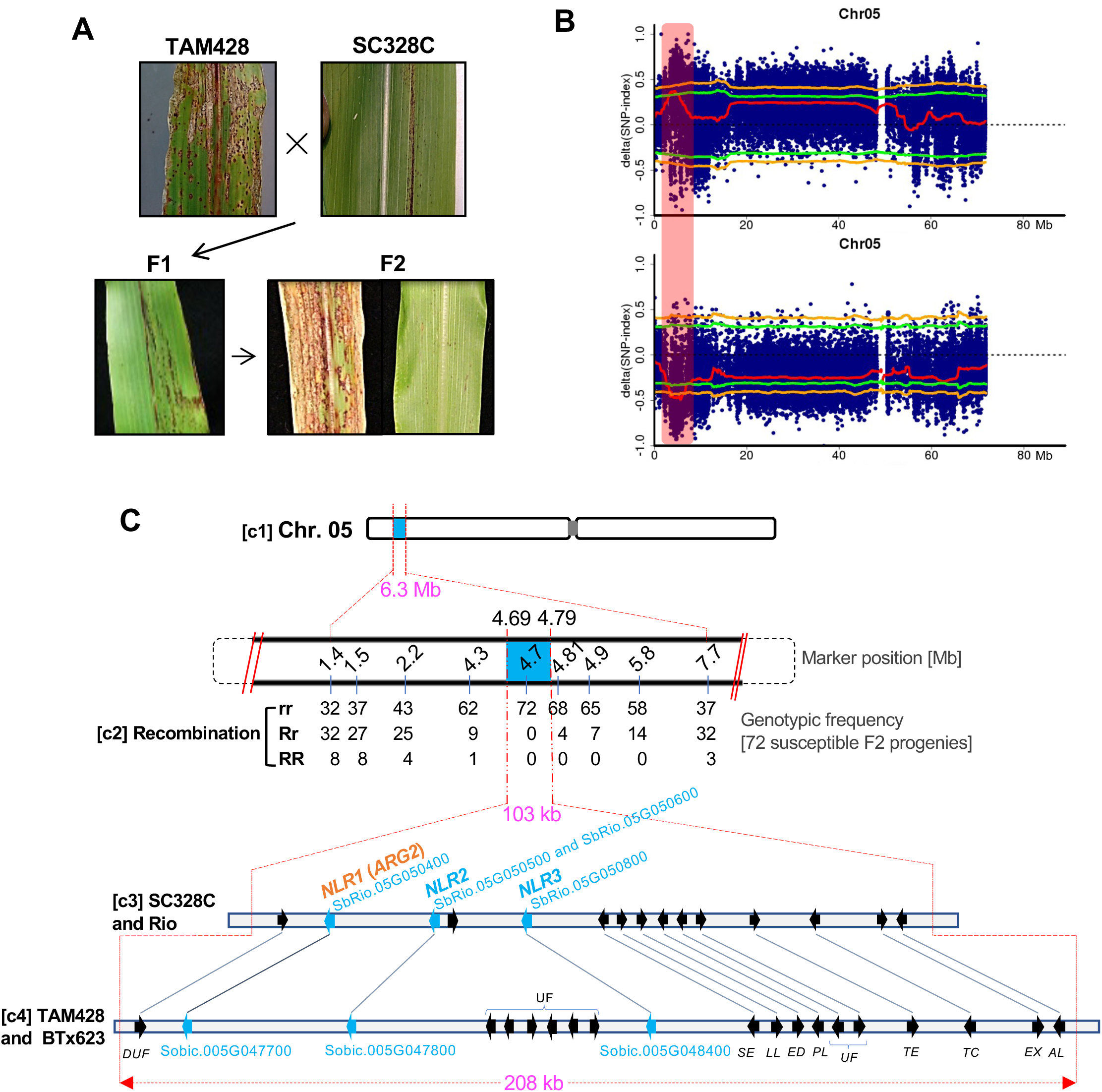
A diagrammatic summary of *ARG2* genetic mapping. **(A)** Disease symptom and resistance response on parents, F_1_ progenies, and F_2_ progenies of the *ARG2* mapping population. **(B)** BSA-seq result showing *ARG2* locus with respect to the resistant (upper) and the susceptible (lower) parental lines. The x-axis is genomic coordinate, and the y-axis is ΔSNP-index estimate. Each blue dot (highly overlapped) represents ΔSNP- index estimate in the 2 Mb sliding window, and the red line is the ΔSNP-index threshold. The green line shows a significance threshold (*p*=0.05) of the ΔSNP-index and the orange line at *p*=0.01. The vertical rectangular bar (light-red shaded) marks the significant *ARG2* mapping region. **(C)** *ARG2* fine mapping. **(c1)** *ARG2* mapping region (6.3 Mb) as identified using BSA-seq. **(c2)** Genetic linkage analysis. The row of numbers (upper panel) are positions of size-markers that were used to generate the genotypic frequencies (**c2**, lower panel). **(c3)** The 103 kb *ARG2* mapping interval defined by the recombination analysis that carries 15 predicted genes in the Rio reference genome. **(c4)** The corresponding genomic interval in the BTx623 reference genome (208 Mb) carries 20 predicted genes. The short arrows in these two *ARG2* genomic intervals show the relative position and orientation of the genes with the homeologs linked using lines. The gene annotations are *DUF*, Domain of unknown function; UF, Unknown function; *NLR, NBS-LRR*; SE, Serine esterase; *LL*, Limkain B Lkap; *ED*, Nad Dependent Epimerase/Dehydratase; *PL*, TRNA-Nucleotidyl Transferase/Poly A polymerase family member; *TE*, TRANSCRIPTION ELONGATION FACTOR B POLYPEPTIDE 3; *TC*, CCR4-NOT Transcription complex subunit; *EX*, Exocyst Subunit EXO70 family protein C1-related. *ARG2* is *NLR1* as identified latter.

To identify the genomic region that carries *ARG2*, composite interval mapping was conducted using BSA-Seq (Takagi et al., 2013). Consistent with the above genetic data, BSA-Seq analysis in F_2_ identified a single genomic region around the proximal end of chromosome 5 that was strongly associated with *ARG2* resistance (Figure 2, B and C). In BSA-Seq, the statistical measure used to determine the causal genomic region of the disease phenotype is a ΔSNP-index (Supplemental Table S2), which is expressed as the difference in the average SNP frequencies between the resistant and the susceptible pooled-progeny genomic samples. ΔSNP-indices were estimated using a 2 Mb sliding window that slides 50 kb at a time. The significant ΔSNP-index estimate at 95% confidence interval (CI) ranged from 0.36–0.50; with the highest threshold estimate of 0.35 at 95% CI and 0.41 at 99% CI. The significant genomic interval was >3.6 Mb in the 95% CI and about 2.8 Mb in the 99% CI. Within the 95% CI, the smallest depth of aligned raw-reads ranged from 24.5 to 28.6 (mean 26.2) and the SNP count ranged 1,067–2,872 (mean, 1999). Based on the SNP index in the BSA-seq, there was no other genomic region that was associated with the resistance in SC328C (Supplemental Figure S2).

### Fine mapping of ARG2 locus

To narrow the *ARG2* mapping interval that is generated using BSA-Seq, genetic linkage analyses was conducted using PCR (polymerase chain reaction) size markers (Figure 2C). *ARG2* is inherited as a dominant allele, and the phenotype of heterozygote F2 plants is resistant, whereas susceptible progenies carry *ARG2* locus in the homozygote state. To reduce the time required for the analysis of the more frequently occurring heterozygote F2 plants, linkage analysis was conducted using 72 susceptible F_2_ progenies, which were homozygous for the susceptible allele of *ARG2*. This analysis resulted in a convergent gradient of allelic frequencies around the *ARG2* locus that narrowed the *ARG2* mapping interval to about 0.51 Mb, which is between the coordinates 4.3 Mb and 4.81 Mb in the Rio reference genome (Figure 2C). Additional genotyping of 336 F_2_ progenies using size markers within this 0.51 Mb interval showed complete co-segregation between *ARG2* phenotypic and genotypic classes, but additional recombinant progeny was not found. After further recombinant analysis, two more recombinants were identified, one at 4.79 Mb in a homozygous resistant progeny and the other at 4.69 Mb in the *ARG2*-NILs. In sum, linkage analyses resulted in a 103 kb *ARG2* mapping interval that is flanked by markers at 4,686 kb and 4,789 kb. Thus, the *ARG2* mapping interval in the Rio reference genome show about a 103 kb difference. The predicted genes *SbRio.05G050200* (4,682 kb) and *SbRio.05G051800* (4,792 kb) flank this *ARG2* mapping interval. In the BTx623 reference genome, these flanking markers are located 208 kb apart (4,492-4,700 kb), and the corresponding flanking genes were *Sobic.005G047300* (at 4,487 kb) and *Sobic.005G049350* (at 4,702 kb). The difference in the *ARG2* mapping interval between Rio and BTx623 is similar between SC328C and TAM428 as well (Figure 2C).

### Identification of a candidate *ARG2* gene

Throughout the *ARG2* mapping interval, the integrated genome view of the parental genome alignment map (BAM files) showed both striking sequence similarities and disparities among the four variants of sorghum: BTx623, TAM428, SC328C, and Rio (Supplemental Figure S3; Supplemental Figure S4). Rio is a sweet sorghum variant that produces high vegetative biomass (Davila-Gomez et al., 2011; Cooper et al., 2019) whereas SC328C has a short stature and produces large panicle. SC328C and Rio, both of which are resistant, carry nearly identical genomic sequences in the 103 kb *ARG2* mapping interval despite their distinct origins (Cooper et al., 2019), and TAM428 and BTx623, both of which are susceptible, carry highly similar genomic sequences in the 208 kb *ARG2* mapping interval although the similarity between resistant lines is higher. Thus, whole genome sequence of the Rio and BTx623 reference genomes were leveraged for the *ARG2* fine mapping.

The entire genomic sequence and all the predicted genes in the *ARG2* mapping interval were examined using the parental sequences and annotated gene functions. The 103 kb *ARG2* mapping interval carries 14 predicted genes (Figure 2C). Four of the 14 predicted genes (*SbRio.05G050400*, *SbRio.05G050500*, *SbRio.05G050600*, *SbRio.05G050800*) encode putative nucleotide-binding site leucine-rich repeat (NB-LRR, NLR) proteins with *N*-terminal coiled-coil (CC) domain, CC-NB-LRR. The remaining genes are identical or nearly identical between the parental genomes and do not show any high impact SNP. These *NLR* genes show high sequence similarity to each other, which caused ambiguous alignments to the reference genomes; parental DNA raw-reads map to more than one of these *NLR* gene regions (Supplemental Figure S3; Supplemental Figure S4).

*ARG2* is inherited as a dominant allele, suggesting that resistance is likely conferred by a functional allele. Accordingly, we first identified the intact candidate genes in the resistant parent. *NLR3* (*SbRio.05G050800*) carries a >9 kb transposon insertion that separates part of the CC domain and the promoter from the rest of the gene and thus is unlikely to be functional (Figure 3). *NLR2* (predicted as two genes: *SbRio.05G050600* and *SbRio.05G050500*) is actually a single *NLR* gene, which can be viewed by comparing the Rio to the BTx623 reference genomes. A single transcript was amplified from Rio and verified by sequencing. In the resistant lines SC328C and Rio, *NLR2* carries a 442 bp duplication in the middle of the gene, which disrupts the ORF and results in mis-annotation of the single gene as two separate genes. Like *NLR3*, this gene is unlikely to be functional. Similarly, comparative genomic (CoGe) analysis of these three highly similar *NLR* genes indicated that two of the three *NLR* genes are disrupted in SC328C and Rio, making *NLR1* (*SbRio.05G050400*) the only viable candidate *ARG2* gene (Figure 3A).

**Figure 3.**
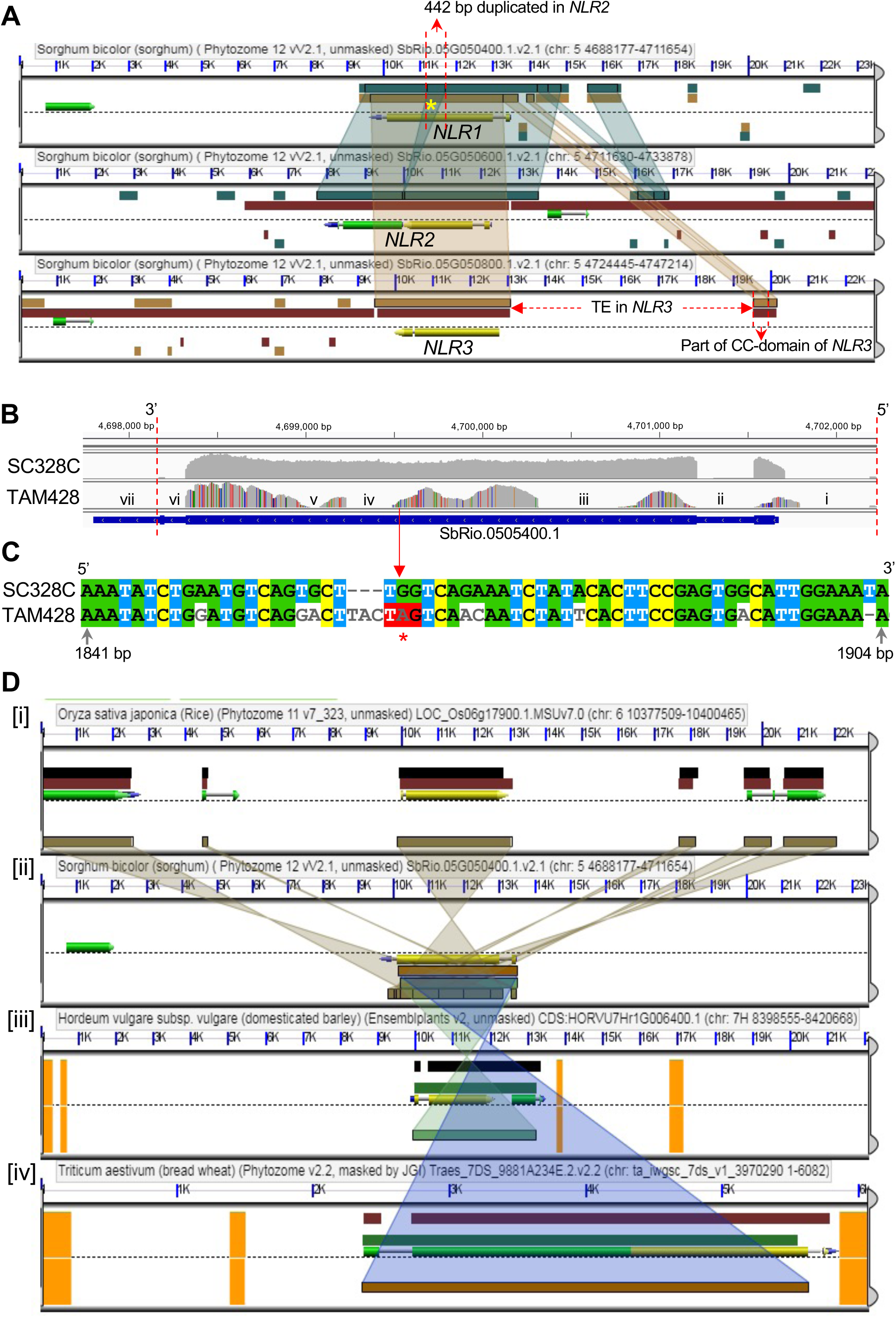
Comparative genomics (CoGe) of *ARG2* duplicate genes and sequence variation of *ARG2* in parental lines. **(A)** CoGe view of the three duplicate *NLR* genes in the *ARG2* mapping interval using the Rio reference genome. The three panels show the three duplicate genes, and the colored connectors between the panels show homologous segments. *NLR2* (in the middle panel) was predicted as two genes (*SbRio.05G050500* and *SbRio.05G050600*) although these two predicted genes are parts of a single gene and code for a single transcript. The dark-green overlapping connector between *NLR1* and *NLR2* (marked with asterisk) shows a 442 bp segment of *NLR1* that is a duplicate in *NLR2*. The bottom panel shows a big transposon insertion (TE in *NLR*3) that disrupted the upstream gene region of *NLR3*. **(B)** The integrated genomic view [IGV] of *NLR1* transcript raw-reads of the parental lines of *ARG2* mapping population that are aligned to the Rio reference genome. These sequences were obtained using wide-Seq. The numbers on top are genomic coordinates of *ARG2*. The vertical distance of the gray area shows the depth of aligned reads, which ranges from 700 to >7,000. The uniformly gray area shows that aligned reads of SC328C are identical to the reference genome, and the SNPs in TAM428 are shown by red, orange, blue, and purple vertical lines in the gray background. The gene regions in the TAM428 panel that does not show aligned reads (no gray area) are [iv] a 62 bp indel and [iii and v] exon segments with dense SNPs that prohibited alignment. The remaining no-gray regions [i, ii, vi] are intronic segments and [vii] part of the 3’UTR that was not included in the sequencing. The gene structure (below the panels) is the predicted *NLR1* gene in the reference genome. **(C)** Aligned coding sequences that carry the premature stop codon in the susceptible parent. **(D)** A snapshot of the top *ARG2* homologs in rice, barley and wheat that were identified using CoGe. The different color connectors between the panels show the homologous segment between genes in the different species. [i] *O. sativa* has a cluster of three highly homologous genes, one of them has gene ID LOC_Os06g17880 (Apoptic ATPase, LRR annotations like ARG2). [iii] *H. vulgare* (barley) homolog has the gene ID HORVU7Hr1G006400. [iv]The homology with top homolog in *T. aestivum* covers 99% peptide segment, gene ID Traes_7DS_9881A234E [RPM1, Apoptic ATPase, NB-ARC, LRR annotations.

Transcript analysis of these three *NLR* genes in the resistant parent supported the above structural genomic analyses (Figure 3, B and C). Transcripts of *NLR1* and *NLR2* were amplified using gene-specific primers in the UTR regions, and these PCR products were used as templates to clone the coding sequences of *NLR1* and *NLR2* into a plasmid vector. There was no transcript amplification corresponding to the *NLR3* gene. *NLR2* has a premature stop codon near the predicted translation start site, and downstream of the 442 bp duplication. Thus, *NLR2* and *NLR3* are disrupted in the resistant parent, and *NLR1* is the only likely *ARG2* candidate gene. *NLR1* is designated *SbRio.05G050400* in the Rio v2.1 and *Sobic.005G047700* in the BTx623 v3.1.1 reference genomes in the phytozome database. Because *NLR1* is intact in the resistant parent, we refer to this as a wild-type allele whereas the gene in the susceptible parent carries a premature stop codon and hence is referred to as a mutant non-functional allele.

### ARG2 cis-regulatory elements carry structural variants

The *ARG2* locus contains a cluster of three duplicate *NLR* genes that have undergone many structural rearrangements including high impact indels (Figure 2C; Figure 3A). In the susceptible lines TAM428 and BTx623, these three duplicate genes are interspersed with several other predicted genes (Figure 2C) within the *ARG2* mapping interval. In contrast to the resistant lines, the susceptible lines do not have the 442 bp duplicate segment in *NLR2* and the large insertion in *NLR3* (Figure 3A). The resistant *ARG2* allele harbors a 480 bp intron in the 5’UTR region, and its transcript carries a 71-codon uORF (Figure 4A). One of the splice sites of this intron has an SNP, C<u>T</u>/CC in the wild-type and C<u>A</u>/CC in TAM428 and BTx623, that eliminates the canonical splice site present in the resistant allele. Nevertheless, this intron carries many SNPs and three small indel sites, and the transcripts in TAM428 and BTx623 have another common splice site in that 480 bp intron, which is TGAT in the wild-type, and T<u>AG</u>GAT in TAM428 and BTx623 (Figure 4A).

**Figure 4.**
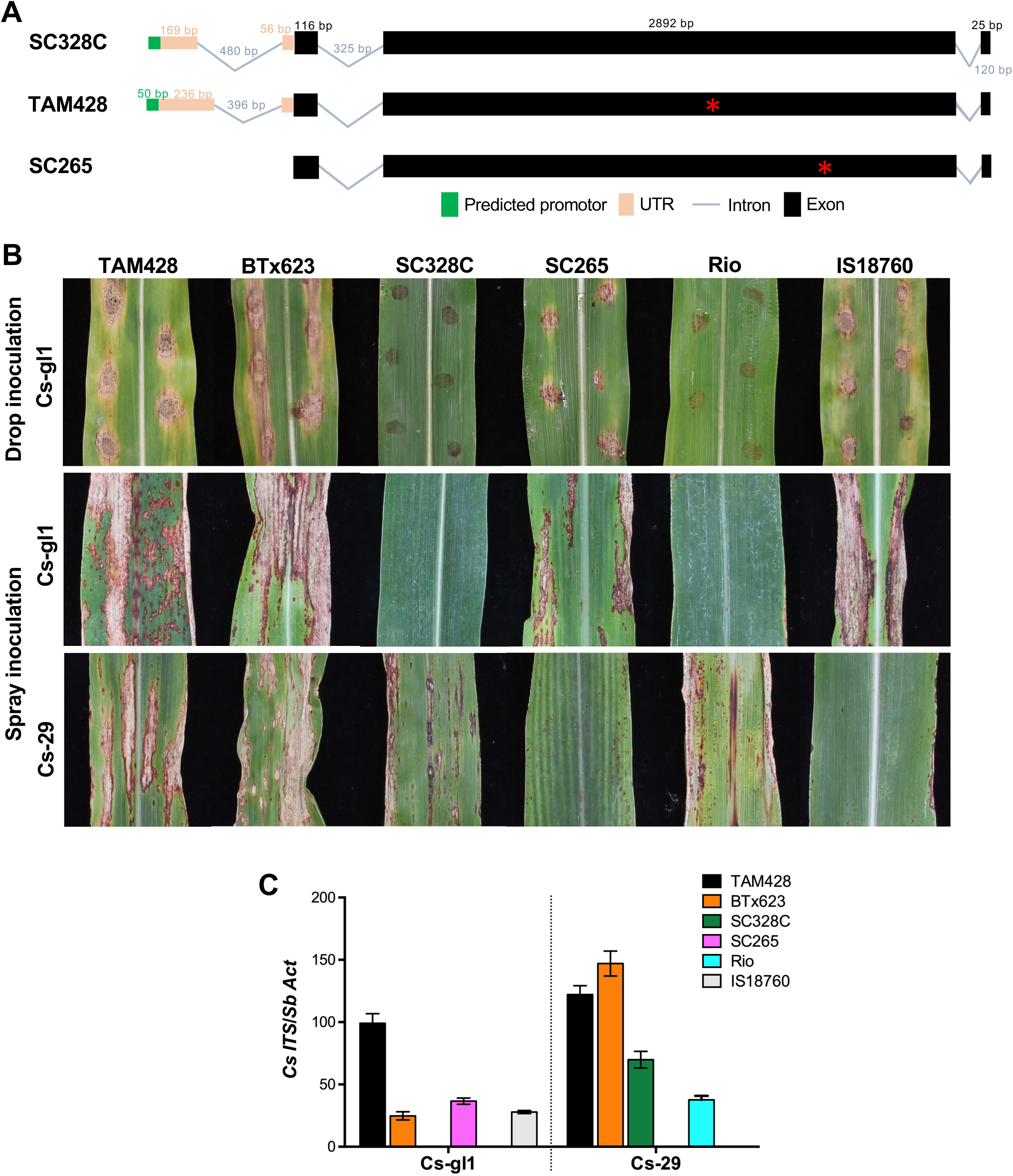
Genomic structure of *ARG2* variants, disease symptoms and pathogen growth. **(A)** *ARG2* domain structure and variations in the upstream cis-regulatory elements of the resistant and susceptible ARG2 alleles. *ARG2* harbors three exons with a complete ORF in SC328C whereas the allele in TAM428 and SC265 carry different premature stop codons. *ARG2* shows structural rearrangement in the 5’UTR between the parental lines of *ARG2* mapping population. **(B)** Sorghum lines that show differential responses to *Colletotrichum sublineola* strains Csgl1 and Cs29. TAM428 and BTx623 are susceptible to both strains of *Cs*. All the images were taken at 7 days after inoculation. **(C)** Growth of *Colletotrichum sublineola* strain on sorghum lines carrying different *ARG2* alleles.

### Validation of ARG2 through independent loss of function alleles

To validate the single *ARG2* candidate gene, *NLR1*, sorghum natural variants were assessed with emphasis on susceptible lines for identification of independent *ARG2* loss of function alleles. In addition to phenotypic evaluations, extensive assessment of *NLR1* was conducted using BAM files of publicly available genomic data. Subsequently, *NLR1* transcripts were amplified from 11 sorghum lines that are susceptible to Csgl1 (SC265, TAM428, BTx642, SC23, IS9830, PQ-434, IS18760, ICSV745, IS8525, Malisor84-7, BTx623) and four resistant sorghum lines (SC328C, Keller, SC237, Rio), and the complete coding sequences of these *NLR1* alleles were sequenced using Wide-Seq. There was no amplification of the *NLR1* transcript and genomic sequence in the susceptible sorghum lines Greenleaf and IS3121 (Supplemental Table S3; Supplemental File S1). The coding sequences in all the 15 sorghum lines were clustered into four similarity groups based on sequence identity, presence of complete ORF and position of the premature stop codon (Supplemental File S2). The four resistant sorghum lines carry identical *ARG2* coding sequences. By contrast, the coding sequences in the susceptible lines were clustered into three types: six lines carry the TAM428 allele, four lines carry the BTx623 allele, and SC265 carries a unique allele. Figure 4B shows the disease phenotype on selected sorghum variants that represent the four different alleles after inoculation with two *Cs* strains that show contrasting virulence. TAM428 and BTx623 show increased disease lesion size and disease symptoms to Csgl1 and Cs29 (Figure 4B).

The IS18760 and SC265 sorghum lines are susceptible to Csgl1 and resistance to Cs29. Conversely, SC328C and Rio were free of disease symptoms in response to Csgl1 after drop and spray inoculations. qPCR using primers on the fungal nuclear ribosomal DNA internal transcribed spacer (ITS) showed no fungal accumulation in SC328C and Rio plants inoculated with Csgl1 (Figure 4C). However, both resistant lines SC328C and Rio showed increased disease symptoms and fungal growth when inoculated with Cs29 (Figure 4, B and C). These data confirmed the robust but race specific resistance in SC328C and Rio to the Csgl1 strain.

*NLR1* in TAM428 and SC265 carries premature stop codons at different positions, and sequence comparisons support an independent divergence of these two alleles (Supplemental Figure S5; Supplemental File S1; Supplemental File S2). As compared to the wild-type *ARG2*, the allele in SC265 carries <8% SNPs whereas the allele in TAM428 carries >10% SNPs. The amino acid sequence differences were also compared based on the shortest ORF, which is 1,848 bp in the TAM428 allele (Supplemental File S3). As compared to the wild-type *ARG2,* in this shortest ORF, the SC265 ARG2 allele carries 4% nonsynonymous codons whereas the allele in TAM428 carries 14% nonsynonymous codons. This ORF in SC265 carries about 15% nonsynonymous codon as compared to the one in TAM428. Thus, the sequence divergence between the susceptible alleles in SC265 and TAM428 is greater than their divergence from the resistance allele. These data show that the mutant allele in the susceptible lines TAM428 and SC265 are distinct because the mutation that inactivated the gene in SC265 must have occurred after the divergence of TAM428, from the resistance alleles in SC328C and Rio. In sum, the *NLR1* allele in SC265 is an independent mutant allele that validates *NLR1* as the functional *ARG2* gene that was defined genetically.

The susceptible BTx623 carries a recessive allele of *ARG2* but retains a complete ORF like the resistance allele in SC328C. The segregation of *ARG2* alleles in the BTx623 x SC328C population also suggested the single locus resistance that is inherited as a dominant allele. The BTx623 *ARG2* allele was examined for sequence differences, as compared to the resistance allele, that could account for the loss of ARG2 resistance function to Csgl1 (Supplemental File S2). Consistent with loss of ARG2 function in BTx623 and TAM428, their *ARG2* sequences are closely related to each other than either is to the resistance allele. In the LRR domain that is crucial for pathogen recognition, as compared to the resistance allele, the allele in BTx623 shows about 19% nonsynonymous amino acid substitutions (Supplemental File S4). This sequence difference in the LRR domain includes two indels (2 and 4 bp) that flank a 45 bp coding sequence resulting in a reading frame shift of 15 amino acids. Together, these data support the loss of *ARG2* resistance in BTx623 to the *Cs* strains Csgl1 and Csgrg to which the allele in SC328C and Rio confers resistance.

The amino acid sequences of ARG2 proteins from SC328C and BTx623 were compared to the HOPZ-ACTIVATED RESISTANCE 1 (ZAR1; template c6j5tC) due to the high amino acid sequence similarity to ARG2 and the availability of solved structure. This comparison revealed that the tryptophan at the 156^th^ amino acid in SC328C is substituted with Serine in BTx623 (Supplemental File S2). A mutation from a conserved tryptophan to alanine at 150^th^ amino acid position of ZAR1 reduces NLR protein oligomerization and consequent cell death, resulting in the increased susceptibility of *Arabidopsis* to *Xanthomonas campestris* pv*. campestris* (Wang et al., 2019). To cover more potential amino acids important for ARG2 function, we aligned sorghum ARG2 from SC283C and BTx623 with top 100 monocot and dicot homologues. The amino acids in ARG2 that are highly conserved in the monocot and dicot ARG2 homologues also show a striking difference between the SC328C and BTx623 alleles (Supplemental Table S4). We found that the conserved amino acids of ARG2 in SC328C are changed in BTx623 at positions 48, 103, 225, 242, 480, and 866 amino acids, respectively (Supplemental Table S4). These conserved amino acid changes imply that the defect of ARG2 function in BTx623. Particularly, Valine (Val) at the 225^th^ amino acid that is highly conserved both in monocots and dicots is retained in the resistant line SC328C but substituted by Leucine (Leu) in the susceptible BTx623 (Supplemental Table S4). Previously, NLR protein autoactivation was observed when Val at this site is substituted with Alanine (van Ooijen et al., 2008).

To further understand differences in ARG2 protein between SC328C and BTx623, their predicted tertiary structures were compared using Phyre2 (Kelley et al., 2015). These structures were compared to the disease resistance RPP13-like protein 4 (ZAR1; template c6j5tC), which suggested that the ARG2 protein in SC328C has a unique structure. First, the beta-sheet structure in BTx623 was replaced by an alpha-helix structure at the NBS region in SC328C (Navy blue color, Supplemental Figure S6A). These changes occurred between the subdomain RNBS-A and Walk B (Supplemental File S2), and the substitution of the highly conserved Val by Leu occurred in this RNBS-A motif. Second, an extra alpha-helix is present between the conserved ARC1 and ARC2 in SC328C (green, Supplemental Figure S6B; Supplemental File S2). The NBS-ARC1-ARC2 structure is important for R protein activation through ATP/ADP binding (Tameling et al., 2002). Third, more alpha-helix was observed around MHD motif (green, Supplemental Figure S6C; Supplemental File S2). We also predicted the ligand binding site of ARG2 using COFACTOR and COACH of I-TASSER software (Roy et al., 2010). This analysis revealed that there is no difference in ATP binding site residues in these two alleles. However, multiple ADP binding site residues between SC328C and BTx623 were altered (Supplemental Figure S6D). The current model suggests that ARG2 protein bound with ADP is in an inactive form in BTx623 (El Kasmi, 2021). Although the impacts of changes in the tertiary structure have not been reported, apparently, there are clear differences between these two alleles.

Like TAM428, BTx623 show increased disease lesion size and disease symptoms to Csgl1 (Figure 4B). qPCR using primers on the fungal nuclear ribosomal DNA internal transcribed spacer (ITS) also showed fungal accumulation in BTx623 but not in the SC328C and Rio plants inoculated with Csgl1 (Figure 4C). However, like BTx623, the resistant lines SC328C and Rio showed increased disease symptoms and fungal growth when inoculated with Cs29 (Figure 4, B and C). These data confirmed the susceptibility of BTx623 to Csgl1 whereas SC328C and Rio show resistance revealing race specific resistance.

### The ARG2 resistance allele localizes on the plasma membrane in *N. benthamiana*

To gain further insight into how ARG2 recognize *Cs*, we investigated the subcellular localization of the resistance allele of ARG2 from SC328C in *N. benthamiana* (Figure 5). We fused enhanced Green Fluorescent Protein (mGFP) to the C-terminus of full-length ARG2 (ARG2:mGFP) and transiently expressed in *N. benthamiana.* The subcellular localization was evaluated at 48 hours following agroinfiltration. Laser scanning confocal microscopy with ARG2:mGFP revealed an even distribution of mGFP fluorescence predominantly on the cell periphery with no observable localization to cytoplasmic strands, suggesting plasma membrane localization (Figure 5). To determine whether ARG2:mGFP localizes on the plasma membrane, we co-expressed ARG2:mGFP with an *Arabidopsis* protein known to localize to the plasma membrane, *At*FLS2:mCherry (Helm et al., 2019). The ARG2:mGFP fluorescence signal co-localized with the *At*FLS2-mCherry fluorescence signal (Figure 5), confirming ARG2:mGFP is indeed localized on the plasma membrane.

**Figure 5.**
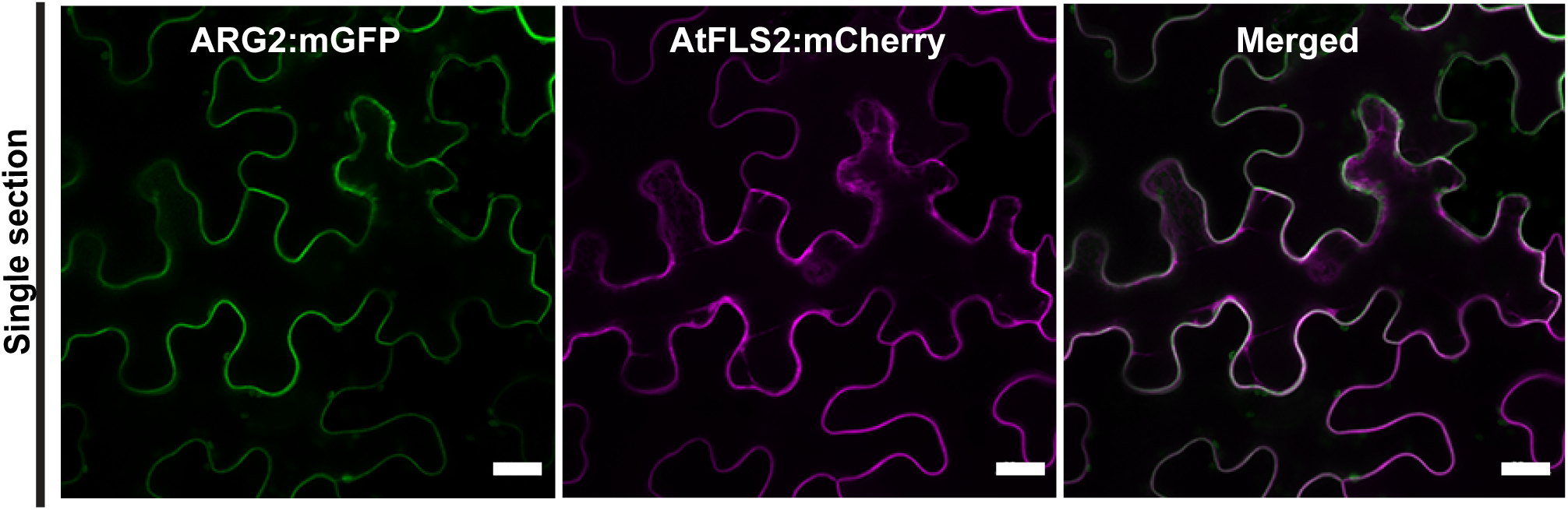
The ARG2 resistance protein, predominately localizes on the plasma membrane. The indicated constructs were transiently co-expressed in *Nicotiana benthamiana* and imaged using laser scanning confocal microscopy 48 hours post-agroinfiltration. mCherry-tagged *Arabidopsis* FLS2 was included as a reference for plasma membrane localization (Helm et al., 2019). The mGFP and mCherry fluorophores were excited at 488nm and 561nm, respectively. mGFP (green) and mCherry (magenta) fluorescence signals were collected between 525 and 550, and at 610nm, respectively. Images shown are single optical sections. The scale bar shown represents 20 μM.

### ARG2 homologs are widely conserved in monocot and dicot plant lineages

Comparative genome analyses revealed that ARG2 has many close homologs in monocot and dicot species. To investigate the relationship between ARG2 in SC328C and ARG2 in BTx623, and other homologous NLR proteins in monocots and dicots, phylogenetic analysis was conducted using amino acid sequences (Supplemental Figure S7). In the phylogenetic tree, the ARG2 homologs in monocots and dicots show distinction. ARG2 protein in SC328C and BTx623 cluster with the other sorghum NLR proteins encoded by the duplicate NLR genes in the ARG2 locus. Despite the widespread homology, the sequence identity of all ARG2 homologs, except the duplicate genes at the *ARG2* locus, were <60%. No ARG2 homolog was identified in the maize genome despite their relatedness, and from other species, *Eragrostis curvula* show the closest ARG2 homolog.

Next, we performed a phylogenetic analysis using amino acid sequences of the NB-ARC domain of ARG2 and functionally characterized NLRs (Kourelis et al., 2021) from diverse plant species. ARG2 clustered in the same clade as the rice resistance genes that confer resistance to the rice blast diseases (Wang et al., 2017) (Supplemental Figure S8). The phylogenetic analysis displayed a clear separation between ARG1, a recently described sorghum NLR (Lee et al., 2022) and ARG2 (Supplemental Figure S8).

### Regulation of *ARG2* gene vary between wild-type and mutant alleles

To shed light on ARG2 function, we examined expression of *ARG2* in response to pathogen inoculation, chitin treatment, and in different plant tissues and growth stages using qRT-PCR (Figure 6). In the resistant line SC328C, the flag leaf showed the highest *ARG2* expression. There was relatively low level of *ARG2* expression in the leaf sheath and the leaf blade at the two earlier sorghum growth stages, and *ARG2* expression was hardly detectable in the stalk and the panicle tissues (Figure 6A). SC328C is resistant to Csgl1 and susceptible to Cs29 whereas BTx623 is susceptible to both strains (Figure 6B). *ARG2* expression was significantly increased at 24 hours post inoculation (hpi) during both resistant and susceptible interactions but at 48 hpi, its expression was strongly induced by Csgl1 in SC328C but was unchanged in response to Cs29. In SC328C, Csgl1 induced expression of *ARG2* was significantly higher than that in response to Cs29. BTx623 showed slightly induced *ARG2* expression in response to both Csgl1 and Cs29 at 48 hpi but was lower than in SC328C. Our results indicate that *ARG2* in SC328C is induced significantly more by Csgl1 to which it confers resistance. The expression of *ARG2* in SC328C was significantly induced in response to chitin treatment than the mock treated samples at 48 hpi (Figure 6C). Chitin is a fungal-derived PAMP that is known to elicit resistance. In contrast, we did not observe induction of the *ARG2* allele in the chitin-treated BTx623 plants, although both Csgl1 and Cs29 can trigger expression of *ARG2* in this line.

**Figure 6.**
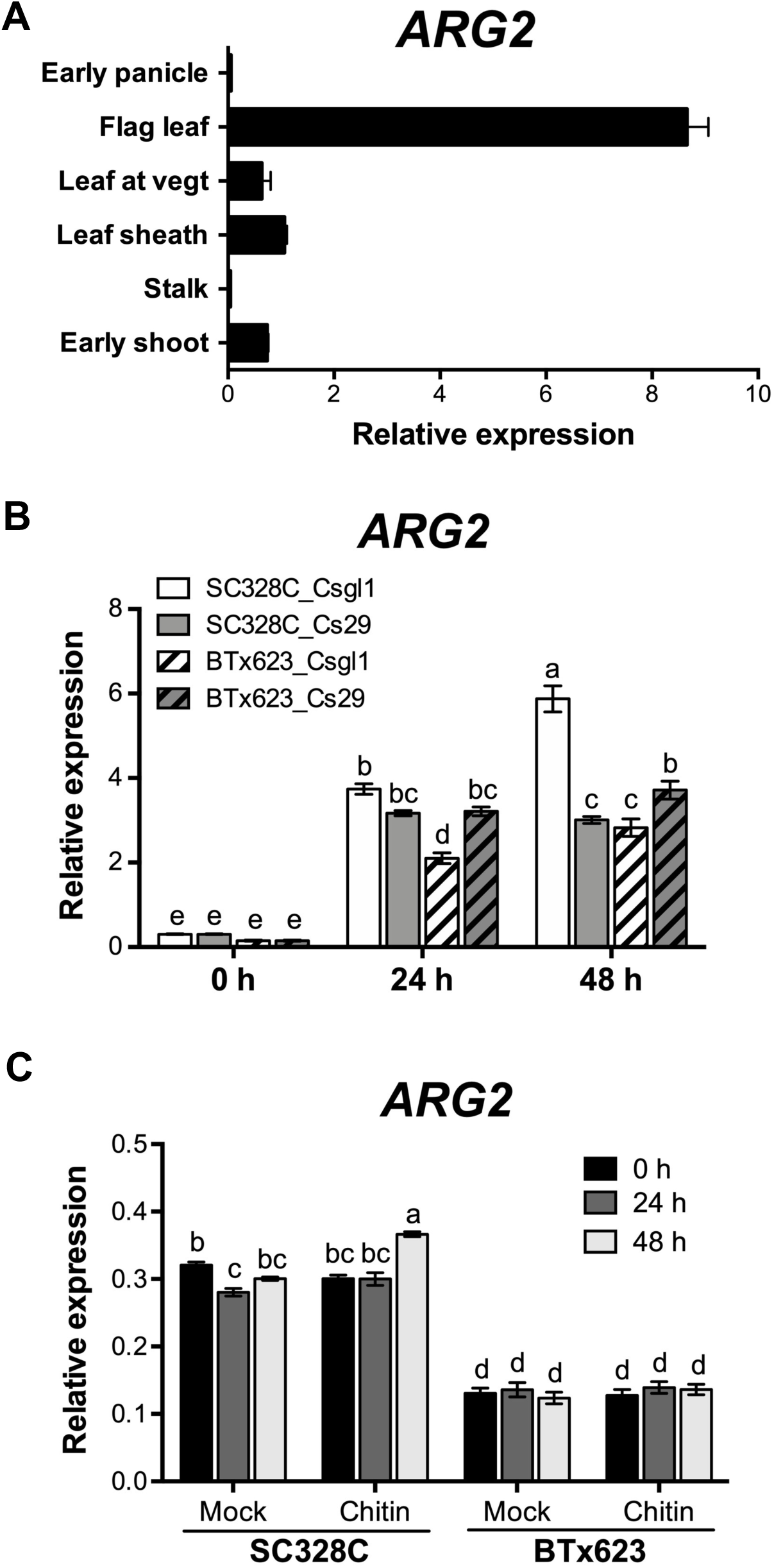
Regulation of *ARG2* gene expression. **(A)** *ARG2* expression in different tissues and growth stages in SC328C. The tissue samples are the entire shoot at 10 days after planting (dap); the stalk, the leaf sheath and the leaf (leaf at vegt) all at 30 dap; the flag leaf at early flowering; and the panicle at early panicle development. Data are mean ± SD from triplicate tissues. Expression level was analyzed using qRT-PCR, and data were normalized by the comparative cycle threshold method with Actin as the internal control. **(B)** *ARG2* expression in response to two different strains of *Colletotrichum sublineola*. SC328C and BTx623 plants were inoculated with *Cs* strains Csgl1 or Cs29, and RNA was extracted at the indicated time points. **(C)** *ARG2* expression in response to chitin. The leaf tissues of SC328C and BTx623 were treated with mock (water) or 2 nM chitin (β-1,4 linked N- acetylglucosamine), and RNA was extracted at the indicated time points. In **(B** and **C)**, expression levels were analyzed using qRT-PCR, and data were normalized by the comparative cycle threshold method with Actin as the internal control. Data show mean ± SE from three independent biological replicates (*P* < 0.05, Tukey-Kramer HSD).

To characterize the functions of ARG2 with minimal interference from differences in the parental genomes, sorghum near-isogenic lines (NILs) were generated by crossing TAM428 and SC328C followed by backcrosses (Figure 7A). The resistant and susceptible NILs inherited the clear-cut difference in disease symptom and pathogen growth of the parental lines (Figure 7B). Expression of eleven defense response genes, selected based on previous studies (Lo et al., 1999; Li et al., 2013), was examined in the NILs using qRT-PCR (Figure 7C). Higher expression of *flavon O-methyltransferase* (*FOMT*), *chalcone synthase 08* (*CHS8*), and *PR10* were associated with resistance at early time points after inoculation (Figure 7C). At later time points, however, the level of expressions of these genes dropped in resistant genotypes whereas the susceptible NILs showed higher levels of expression, which is associated with elevated pathogen growth. Heat shock 90, chalcone synthase 5, pathogenesis related thaumatin-like protein, DNAJ heat shock protein, mitogen activated protein kinase, prolyl 4- hydroxylase, glutathione S-transferase 4, and a defensin homolog GEM-like protein 1 (Li et al., 2013; Mishra et al., 2017) did not show difference in gene expression between the NILs (data not shown). These results suggest increased expression of a subset of sorghum immune response genes in the presence of ARG2.

**Figure 7.**
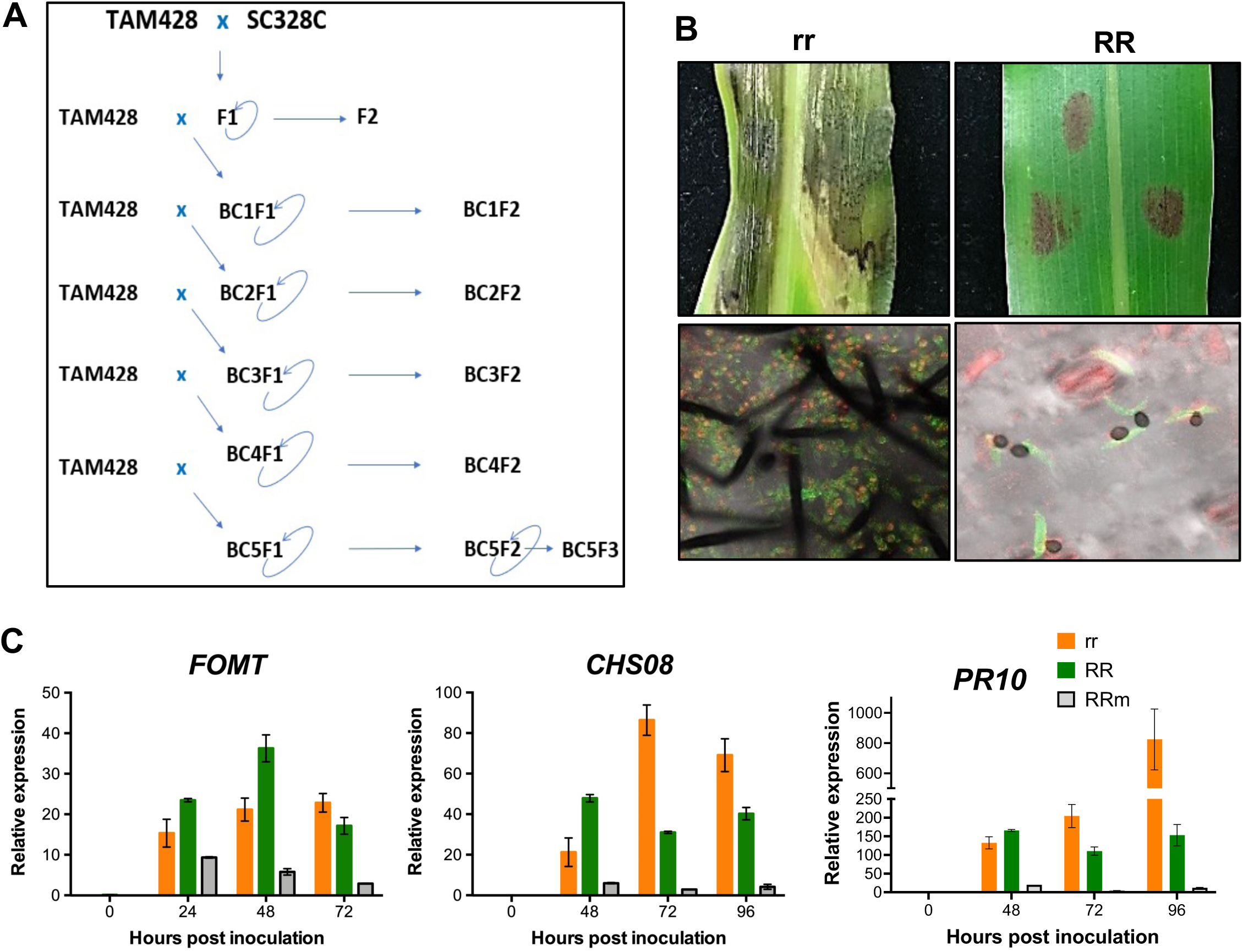
Defense gene expression in near-isogenic lines of sorghum that differ in the *ARG2* locus. **(A)** Schematics showing the steps used to generate the near-isogenic lines (NILs). The NILs carry the recessive (susceptible) *arg2* allele in TAM428 (*arg2/arg2*, *rr*) and dominant (resistance) *ARG2* allele (*ARG2/ARG2*, *RR*). **(B)** Disease symptom and fungal growth in resistant NILs (*RR*) and susceptible NILs (*rr*). **(C)** Expression of three defense response genes in the resistant NILs (*RR*) and the susceptible NILs (*rr*) after pathogen or mock inoculation. *FOMT*, Flavon-O methyltransferase; *CHS8*, Chalcone synthase 08; *PR10*, Pathogenesis related gene 10; hpi, hours post inoculation. Fungal growth on the inoculated tissue was examined using a confocal microscope at 6 days post inoculation (dpi). WGA-AF488 stains fungal hyphae green, and propidium iodide stains dead host cell red. The image was taken at 10 dpi in the detached-leaf disease assay.

### Metabolite profiling data support *ARG2* resistance function

To gain insight into pathogen induced accumulation of secondary metabolites that may contribute to ARG2-mediated resistance, untargeted metabolite profiling was carried out using tissue samples from pathogen and mock inoculated *ARG2* NILs. A total of 2,916 preprocessed metabolite mass-features were obtained: 1,356 in the negative ionization mode and 1,560 in the positive ionization mode. Regardless of genotypic difference, both in the resistant and susceptible NILs, many mass-features were detected only after pathogen inoculation. However, none of these mass-features were detected exclusively in *Cs* inoculated resistant NILs. A total of 474 mass-features showed significant (*p*<0.05) difference between the resistant and the susceptible NILs after *Cs* inoculation (Supplemental Table S5A). Only 57 of these mass-features showed higher accumulation in the resistant NILs, and the rest 417 mass-features showed higher accumulation in the susceptible NILs (Supplemental Table S5A).

Mass spectrometric (MS) “peak to pathways” analysis using the MetPA pipeline showed a total of 86 metabolite hits in 43 pathways (Supplemental Table S5B). Most of these metabolite hits show the mass-features carry abducts like H_2_O or a carbonate salt in addition to ionization feature (Supplemental Table S5C). Six relatively highly enriched pathways are shown in Supplemental Figure S9A. Zeatin and anthocyanin metabolisms showed the highest score upregulation in the *Cs* inoculated resistant NILs followed by mock inoculated resistant NILs. In contrast, *Cs* inoculated resistant NILs showed downregulation of metabolites that are involved in lysine, tryptophan, and arginine and proline metabolisms. Porphyrin and chlorophyll metabolism were downregulated only in *Cs* inoculated resistant NILs.

### ARG2 resistance is not associated with growth tradeoff

The common genomic background in the *ARG2* NILs was leveraged to examine the impact of the *ARG2* locus on plant growth under controlled growth conditions in the absence of the pathogen (Figure 8; Supplemental Figure S9, B-F). Lower leaves of SC328C turn brownish at the vegetative growth stage and part of its panicle remains covered by the leaf sheath (Supplemental Figure S10, A and B). Significantly higher panicle and total dry matter weight were obtained in the true breeding resistant ARG2 NILs as compared to the NILs that carry the recessive homozygote and heterozygote genotypes (Figure 8A). Total leaf area and photochemical reflectance index and nitrogen reflectance index at 97 and 119 days after planting revealed significant differences between the NILs (Figure 8, B-D). In the true breeding resistant NILs, the shoot growth indicator morphometric and color related traits (Vyska et al., 2016; Zhu et al., 2021) and top-plant-surface and side-average-convex-hull were also higher in most plant growth stages (Figure 8, E and F). Further, anthocyanin reflectance index and plant senescence reflectance index differed between the NILs (Supplemental Figure S9, C and D). Some color (hue) related traits showed relatively higher indices in the true breeding resistant NILs in the vegetative stage that is reversed in the reproductive stage (Supplemental Figure S9E) suggesting that traits relatedness may vary with growth stage and a specific growth stage may not sufficiently explain a major aspect of the differences among the resistance and the susceptible NILs. Nitrogen reflectance index showed negative correlation with most of these plant growth, anthocyanin, and senescence traits (Supplemental Figure S9F).

**Figure 8.**
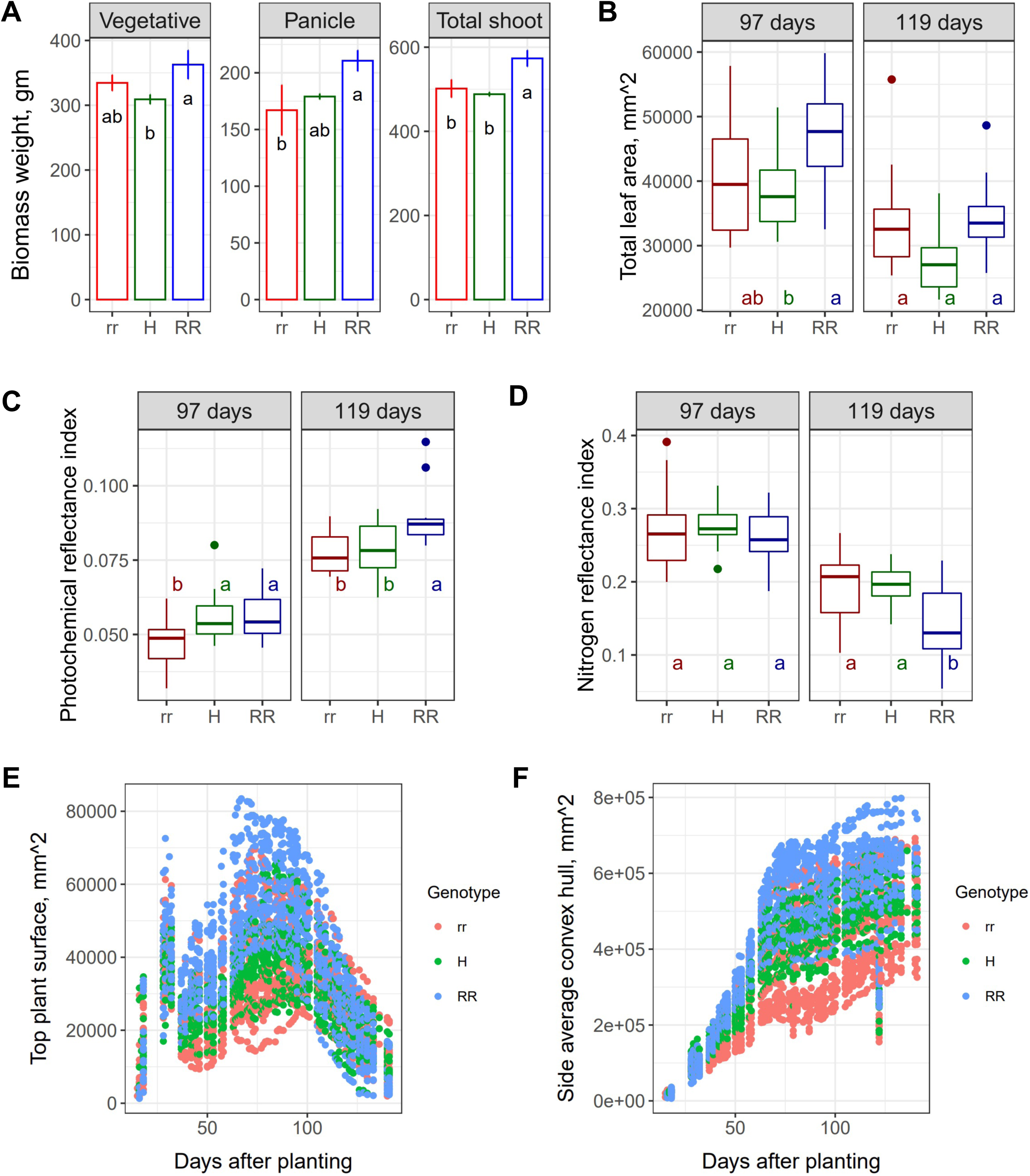
Effect of ARG2 on plant growth in the absence of disease in a controlled environment phenotyping facility. **(A)** Shoot dry matter weight (gm) among *ARG2* genotypes. rr, homozygote recessive; H, heterozygote; RR, homozygote dominant. Shoot dry matter is higher (*p*<0.002) in RR; Panicle dry matter is lower in rr (p<0.02); and vegetative shoot dry matter is lower (*p*<0.015) in the heterozygote, H. **(B)** Total leaf area and **(C)** photochemical reflectance and **(D)** nitrogen reflectance indices at flowering time (97 days after planting, dap) and grain filling stage (119 dap), based on hyperspectral image. Estimates of **(E)** Top plant surface and **(F)** side average convex hull throughout the growth period. rr, homozygote recessive; H, heterozygote; RR, homozygote dominant genotypes. **(B-F)** Estimate of physiological and morphometric traits at flowering time (97 dap – days after planting) and grain filling stage (119 dap) based on red-green-blue (RGB) and hyperspectral image data throughout the growth period. ANOVA and Tukey HSD functions were used to test statistical significance of differences among genotypes.

## Discussion

Here we report the identification and characterization of a sorghum *ANTHRACNOSE DISEASE RESISTANCE GENE2* (*ARG2*), encoding a nucleotide-binding leucine-rich repeat (NLR) protein. *ARG2* was identified based on a biparental mapping population developed by crossing the *Cs* resistant sorghum line SC328C and the susceptible line TAM428. The resistance parent showed complete resistance to the *C. sublineola* strains Csgl1 and Csgrg but susceptibility to other strains. SC328C is one of the sorghum differential cultivars used for race identification (Moore et al., 2008; Prom et al., 2012). However, until now, a specific resistance locus in SC328C against any *Cs* strain had not been identified. Correlation between multiple and distinct *ARG2* alleles and the corresponding host response validated the identity of the *ARG2* gene. For instance, the *ARG2* locus is identical in the sweet sorghum Rio and the dwarf grain sorghum SC328C, consistent with their mutual resistance to Csgl1 and Csgrg but despite major differences in the genetic backgrounds of these genotypes. In addition, multiple loss of functional alleles, including those in the susceptible TAM428 and SC265 lines, that do not encode the predicted ORF support the conclusion that loss of ARG2 is responsible for a loss of resistance. Previously, a minor effect anthracnose resistance QTL was mapped in the genomic region containing the *ARG2* locus based on the susceptible BTx623 and resistant SC748-5 biparental mapping population. Resistance in that case was determined using fungal strains that are different from those in the current study (Burrell et al., 2015). BTx623 and SC748-5 both carry the same susceptible *arg2* allele, and hence the QTL in SC748-5 likely defined a different locus. Interestingly, *ARG2* mediated resistance maintains effectiveness up to 38°C.

Surprisingly, despite the reported growth and defense trade-off associated with resistance genes (Vyska et al., 2016), with or without disease, the NILs that carry the resistance *ARG2* allele grew better than the NILs that do not carry the resistance allele. Each of these observations suggests that the trade-off between growth and disease resistance is ameliorated in this case, making this gene a particularly useful one for introgression.

*ARG2* is found within a complex locus that contains *NLR* genes that have both copy number and structural variations. The *ARG2* locus contains >4 predicted *NLR* genes in the genome of the sorghum line RTx430, and three copies in BTx623 and Rio genomes. Despite the complex nature of the *ARG2* locus, the premature stop codon in *NLR2* and the absence of a transcript from *NLR3* reduced the genomic complexity and allowed us to determine that *ARG2* is the causal gene for the disease resistance. This is in addition to multiple independent *ARG2* recessive alleles that demonstrates loss of resistance. Comparative studies revealed >17,000 copy number variants in sorghum (Zheng et al., 2011) and hundreds of genome specific loci in rice (Schatz et al., 2014).

Support for the role of ARG2 in disease resistance is provided by independent *ARG2* alleles in distinct sorghum lines. Further, *ARG2* expression shows differences between the resistant SC328C and the susceptible BTx623 lines. Thus, multiple lines of evidence suggest functional differences between ARG2 from SC328C and BTx623. Given that, the *ARG2* allele in BTx623 may be useful for a broader understanding of the mechanism of NLR action and evolution of NLR resistance and evolution of NLR specificity. Interestingly, although the *ARG2* allele in BTx623 has an intact ORF, it does not confer resistance to the *Cs* strains Csgl1 and Csgrg. There is, however, enormous sequence divergence relative to the resistant *ARG2* allele in SC328C. Multiple differences in conserved and critical amino acid residues and in predicted structures of ARG2 proteins from SC328C and BTx623 may account for functional differences between the two alleles. The LRR-domain of the BTx623 ARG2 allele shows 19% amino acid sequence divergence relative to the SC328C *ARG2* allele, which may have caused loss of *Cs*-effector recognition from strains avirulent on SC328C. In addition, multiple ADP binding site residues show differences between SC328C and BTx623 and such differences are known to affect function of NLRs (Takken et al., 2006; Williams et al., 2011). The most critical differences include changes that disrupt NLR oligomerization and thus do not trigger cell death, resulting in disease susceptibility (Wang et al., 2020). In sorghum, a high proportion of large effect SNPs are in the LRR domain (Zheng et al., 2011). To our knowledge, there is no *Cs* strain to which BTx623 confers resistance. Although BTx623 is susceptible to all five *Cs* strains tested in the current study and to many more strains in earlier studies (Prom et al., 2012; Klein et al., 2017), the presence of a complete ORF suggests that *ARG2* in BTx623 may confer resistance to other strains (Panchy et al., 2016).

Sequence comparison using CoGe (Figure 4D) and BLASTp identified monocot and dicot homologs of ARG2. However, the sequence similarities of all the homologs, except that of the *NLR* genes in the *ARG2* locus, were less than 60%. This is not unusual among NLR genes that are subject to frequent positive selection (Van de Weyer et al., 2019). A phylogenetic analysis of functionally characterized NLRs including ARG2 reveals that ARG2 clusters with rice Pti2, Pti9, PigmR/PigmS, and Piz-t resistance genes, which confer resistance to rice blast (Wang et al., 2017). Finally, NLR-mediated resistance is known to be modulated by temperature (Bao et al., 2014). The stability of ARG2 at high temperature and absence of resistance-growth tradeoff suggests that ARG2 confers a particularly useful form of resistance. *ARG2* confers race-specific resistance, thus, identification of the *Cs* effector recognized by ARG2 will advance molecular dissection of resistance to *Cs*.

## Materials and Methods

### Plant materials and fungal culture

To identify resistant lines, a collection of sorghum lines was screened for resistance to *Colletotrichum sublineola* (*Cs*) strains Csgl1, Csgl2, Csgrg, Cs27, and Cs29. Csgl1 and Csgl2 were collected in Indiana (USA) (Jamil and Nicholson, 1987). Csgrg was a recent collection from Tiffon, Georgia (USA), and Cs29 and Cs27 were from Western Ethiopia. The fungal strains were cultured on potato dextrose agar (PDA, BD Difco™). Fungal culture, spore collection and disease assays were conducted as described recently (Lee et al., 2022).

### DNA extraction and sequencing

Genomic DNA was extracted using the DNeasy plant mini kit (QIAGEN). The quality and concentration of the DNA samples were determined on gel electrophoresis and quantified using a nanodrop. The DNA samples from individual F_2_ plants were then used to constitute two pooled DNA samples, one for the susceptible group and the other for the resistant group. Equal amounts of DNA from each of the 50 resistant and the 50 susceptible F_2_ progenies of the cross between TAM428 x SC328C was constituted as pooled DNA samples for sequencing. The pooled samples were sequenced at 25x coverage using Illumina HiSeq 2500.

### BSA-seq and genetic linkage analysis

Bulk-segregant analysis of the genomic DNA raw-reads (BSA-Seq) was carried out using the QTL-seq pipeline (Takagi et al., 2013) to identify the genomic region that confers resistance in SC328C to Csgl1. This pipeline uses the pooled-genomic raw-reads to locate the genomic regions that are significantly associated with the resistance. All the default statistical parameters as specified in the pipeline were used. The BTx623 and the sweet sorghum Rio reference genomes were used to align the corresponding raw-reads of the pooled samples. The basis of this analysis is the difference between the average frequency of SNPs in the pooled samples (ΔSNP-index) that are inherited from the two parental lines. In the genomic regions that are associated with the disease phenotype, the allelic frequency in the F_2_ pooled-progeny samples is expected to be significantly different from random. Thus, ΔSNP-indices that result from the random assortment of alleles approximate zero whereas ΔSNP-indices around the genomic regions that are significantly associated with the disease phenotype are different from zero and are observed as peaks in the BSA-Seq result.

The detailed statistics of BSA-seq provide a better picture of the potential genomic interval of the candidate locus around the peaks. SNP-index of the resistant and the susceptible pooled samples is estimated as the average frequency of all identified SNPs in the two Mb and the four Mb sliding windows. The windows slid 50 kb between successive SNP-index estimates, and majority of the genomic region overlaps between successive sliding windows. Thus, depending on the density and distribution of the SNPs, differences between adjacent ΔSNP-indices arise from both the dropped 50 kb sequence of the preceding window and the subsequently added 50 kb sequence, and adjacent ΔSNP-indices are estimated from highly overlapping SNPs. This nature of the BSA-Seq results in a large genomic region that harbors the candidate resistance genes.

DNA size-markers were designed in the BSA-seq peak genomic region to narrow the wider genomic interval of the *ARG2* locus. The binary alignment map (BAM files) of the parental and the pooled-progeny genomes were examined using the integrated genome viewer (IGV) software to identify insertion-deletion (indel) sites to design PCR primers (Supplemental Table S5). Accordingly, many polymorphic DNA markers between the parental genomes were designed. The frequency of mismatch between the disease phenotypes and the parental DNA marker-type is expected to increase with distance from the *ARG2* locus. Thus, to generate a strong gradient of allelic frequencies, markers were designed beyond the peak genomic region at which the ΔSNP-indices dropped off to random recombination.

To design primers, once a potential-indel site was identified on IGV, the preferred primer annealing sites were picked manually to avoid SNPs between the parental genomes that may create a difference in the PCR efficiency due to the parental origins of templates. This manual method enabled the design of optimal indel-to-amplicon size proportion as well, which eased the gel electrophoresis separation of bands. The Rio reference genome was not integrated into the NCBI database. Therefore, the specificity of primer pairs was examined using the blast-tool in the phytozome database (https://phytozome-next.jgi.doe.gov/). The sequences from the desired annealing sites were copied from IGV and optimized as primers using the NCBI primer-designing tool. Finally, the specificity of primers was verified using amplicons from the parental lines prior to use for recombination analysis. The marker size determined the parental origin of amplified genomic segments in the progenies based on DNA size differences gel electrophoresis.

The F_3_ families of recombinant F_2_ plants were evaluated to ensure the true-to-type disease phenotype. Genetic linkage analysis was considered complete when the marker-type in a recombinant F_2_ plant was reconciled to the disease phenotype in the respective F_3_ progenies. This cycle of linkage analysis was carried out several times by designing a new set of polymorphic markers.

### Fine mapping

Once the number of candidate genes was reduced, the fine mapping employed comparative and functional analytical tools. Comparative genomics (CoGe) was used to examine aligned BAM files of many sorghum lines. Functional domains were examined in the reference genomes and NCBI, SMART protein and CoGe databases. Targeted sequencing (Wide-Seq) was carried out using PCR products from genomic DNA and cDNA to further examine SNPs, open reading frame (ORF) and functional domain of the likely candidate genes in the *ARG2* mapping interval between the parental lines.

### Generation of *ARG2* near-isogenic lines

Near-isogenic lines (NILs) that differ in the *ARG2* locus were developed through repeated backcrosses to the BC_5_F_3_ generation using the susceptible parental line as a recurrent mother plant. The controlled pollination in the backcrossing scheme was performed using the plastic-bagging method (Laxman, 1997; Lambright, 2019). Because the pollen source is a heterozygote, the subsequent backcross progenies are expected to carry the heterozygote or the recessive homozygote *ARG2*. There was no molecular marker to track the *ARG2* locus when the NILs were developed, and the F1 progenies of the backcross generations were identified primarily using their disease phenotype. This BCF_1_ screening method necessitated the evaluation of BCF2 progenies of each backcross generation to avoid disease escape rather than a resistant BCF1. The BCF_1_ progenies of all the backcross generations that carry the heterozygote genotype at the *ARG2* locus are expected to show the 3:1 phenotypic ratio at BCF_2_.

### Characterization of plant growth

*ARG2* NILs were grown in Purdue University Ag Alumni Seed Phenotyping Facility (AAPF) growth chamber (120 μmols/m^2^/s and 65% relative humidity) to determine the effect of *ARG2* on growth and physiological status of the plants in the absence of disease. Twenty plants each form the true-to-type recessive, or dominant genotypes and sixteen heterozygote NILs were used to examine differences throughout the growth period. The plants were supplied with minimal fertilizer to avoid compensatory growth, which may arise due to a higher amount of fertilizer and shadow any difference among NILs. RGB (Red-Green-Blue) and hyperspectral images were captured once a day (sometimes once in a few to several days); a total of 66 times from the 16^th^ day after planting to physiological maturity. Shoot dry matter was examined among a subset of well-balanced group of genotypes for analysis of dry matter variance (ANOVA). The dry matter analysis was carried out in three replications, and each replication was represented by three plants, a single plant from each of three BC_5_F_3_ families. Plants in each of these families were raised from a single *ARG2* BC_5_F_2_ heterozygote plant.

### Transient expression assays and Laser scanning confocal microscopy

To determine the subcellular localization of ARG2, full-length *ARG2* coding sequence amplified from SC328C and cloned by Gateway (Invitrogen) recombination reactions into the pEarleyGate103 vector, upstream of the mGFP5 under the control of the CaMV 35S promoter. Transient protein expression assays were performed as previously described (Helm et al., 2019). Bacterial suspensions were mixed in equal ratios (1:1) and infiltrated into the 3-week-old *Nb* leaves. Confocal laser scanning microscopy was performed at 48 hours following agroinfiltration using a Zeiss LSM880 upright confocal microscope. Leaf sections were examined on abaxial surface, and at least two different leaves from two different plants were imaged. mGFP protein fusions were excited using a 488nm argon laser and detected between 525nm and 550nm. mCherry fusions were excited using a 561nm helium-neon laser and detected at 610nm. Images were collected on the Zeiss confocal and processed using the Zeiss Zen Blue Lite program (Carl Zeiss Microscopy, USA).

### Metabolite analysis

Ground tissue samples from the susceptible and the resistant *ARG2* NILs at 48 hours after inoculation that were collected for the gene expression analysis, were used for metabolite analysis. Equal weights of the ground tissue from the three biological replicates were aliquoted to Eppendorf tubes for metabolite extraction using 50% methanol and resuspend by vortex. The samples were then incubated at 65°C for 2 hours by vortexing every 30 minutes, followed by centrifugation at 13,000 rpm. The supernatant was transferred to a fresh tube leaving the residue. Each extract was dried for 24 hours using a SpeedVac. Each sample was re-dissolved in 50% methanol and sonicated for 15 minutes and centrifuged at 13,000 rpm for 10 min. The supernatant was transferred to a new tube. The metabolite samples were kept at -20°C until run on HPLC-MS.

Untargeted metabolite data was generated using the HPLC-MS platform in the negative and the positive ionization modes. A thorough manual peak-integration of the metabolite mass-features was carried out on Agilent’s Mass Profiler Professional, and the conversion of peaks to mass-features was performed on the same platform. The MetPA analytical platform was used to visualize the summary statistics, to identify metabolites that showed significantly different concentrations among groups and to generate the input datasets for pathway analysis. The MetPA platform (www.metaboanalyst.ca) was used for pathway analysis using several metabolite databases including the rice and the Arabidopsis KEGG database (Kanehisa and Goto, 2000).

## Author contributions

D.B.M. conducted most of the experiments including the development of experimental population, genetic studies, ARG2 mapping and gene identification, characterization of the ARG2 locus, gene expression and metabolite analyses, and image-based plant growth studies. S.L. conducted identification of ARG2 and cloning of candidate NLRs, generating vector constructs, disease assays and fungal growth analysis, gene expression, and phylogenetic analyses. A.A., conducted bioinformatic and genetic analyses. C-J.L. conducted the tertiary structure predictions. A.M.S conducted image-based data processing for the analysis of growth traits. M.H. conducted the localization experiment. D.B.M., S.L., C-J.L., A.A., M.H., D.L., and T.M. designed the research, analyzed the data, and wrote the paper. T.M, oversaw the project.

## Acknowledgments

This study was made possible through funding by the Feed the Future Innovation Lab for Collaborative Research on Sorghum and Millet through grants from American People provided to the United States Agency for International Development (USAID) under cooperative agreement No. AID-OAA-A-13-00047. The contents are the sole responsibility of the authors and do not necessarily reflect the views of USAID or the United States Government. We also acknowledge grant from NSF (IOS-1916893) to TM. We also thank the Purdue University Imaging Facility for access to the Zeiss LSM880 upright confocal microscope. We would also like to thank Rachel Hiles (Purdue University) for technical assistance. This research was funded, in part, by the United States Department of Agriculture, Agriculture Research Service (USDA-ARS) research project 5020-21220-019-00D. The funding body had no role in designing the experiments, collecting the data, or writing the manuscript. Mention of trade names or commercial products in this publication is solely for the purpose of providing specific information and does not imply recommendation or endorsements by the United States Department of Agriculture. USDA is an equal opportunity provider and employer.

## Supporting Information

**Supplemental Figure S1. Fungal growth and anthracnose disease symptom at high temperature.**

(A) Fungal growth in resistant (Sc328) and the susceptible lines (BTx623) differing in the ARG2 gene as revealed by trypan blue staining of spray inoculated tissues. The dark stain shows fungal hyphae. (B) Disease symptoms in SC328C at high temperature (38°C), and (C) Disease symptoms on the resistant and the susceptible ARG2 near-isogenic lines at 38°C. The susceptible TAM428 genotype was killed by the high temperature. The resistance in SC328C to Csgl1 was compared to the widely known resistant line SC748. The appearance of disease symptoms at 38°C was delayed relative to the disease assays in laboratory (22-23°C) and greenhouse (30-32°C). The image was taken at 15 dpi.

**Supplemental Figure S2. BSA-seq analyses showing the mapping of the ARG2 locus.**

Pooled DNA from resistant and susceptible F3 plants was sequenced and used for the BSA-seq analyses with respect to (A) resistant parent, and (B) susceptible parent. The x-axis is genomic coordinate, and the y-axis is ΔSNP-index estimate. Each blue dot, highly overlapped, represents ΔSNP-index estimate in the 2 Mb sliding window, and the red line is the ΔSNP-index threshold. The green line shows a significance threshold (*p*=0.05) of the ΔSNP-index and the orange line at *p*=0.01.

**Supplemental Figure S3. Aligned BAM files as examined in integrated genome viewer (IGV).**

SC328C, TAM428 and BTx623 genome raw-reads binary alignment map, BAM, files aligned to the Rio reference genome. The lower panel of each sorghum line shows aligned reads and the coverage, and the upper panel shows the frequency of aligned raw-reads. The upper image shows the alignment at NLR1 (*ARG2*), and the lower image shows alignment at NLR2 and NLR3. During mapping, reads that found perfect match align without ambiguity and even highly similar reads from duplicate genes align without ambiguity, and hence the BAM coverage in IGV appears uniformly gray, as in SC328C. These uniformly gray shaded areas in the coverage panels show no SNP. ARG2 sequence in SC328C is identical to Rio whereas enormous variation exists with TAM428 and BTx623 that resulted in ambiguous alignment of reads as shown by red, blue, green, and brown vertical lines (genomic coordinates) and some positions with no aligned reads. Each of these colors represent one of the four nucleotides as SNPs. Multiple color in a single coordinate (vertical line) of a coverage panel (blue and red, for example) shows reads that belong to multiple duplicate genes.

**Supplemental Figure S4. General mapping pattern of sorghum variants in the ARG2 genomic region covering 80 kb.**

General mapping pattern of 16 representative sorghum lines in the *ARG2* locus that cover 22 kb genomic region. The 16 panels show mapping statistics of the 16 variants that are mapped to the Rio reference genome. The top panel belongs to SC328C (the resistant parent), and all the rest belong to mutant *arg2* variants. The *NLR1* (*ARG2*), and the non-functional NLRs are shown.

**Supplemental Figure S5. The evolutionary relationship of ARG2 among 15 sorghum lines.**

Circles represent resistant and diamonds represent susceptible lines to Csgl1. Initial tree(s) for the heuristic search were obtained automatically by applying Neighbor-Join and BioNJ algorithms to a matrix of pairwise distances estimated using the JTT model, and then selecting the topology with superior log likelihood value. The tree is drawn to scale, with branch lengths measured in the number of substitutions per site. There are a total of 1,011 positions in the final dataset. Evolutionary analysis was conducted in MEGA X. Bootstrap values are given at the node as a percentage of 1000 replicates.

**Supplemental Figure S6. Prediction of 3D structure of resistant and susceptible ARG2 alleles.**

(A-C) The tertiary structure of ARG2 predicted based on the template c6j5tC (RPP13-like protein 4). The ARG2 structure of SC328C and BTx623 was predicted by Phyre2 (Kelley et al., 2015).The specific regions of ARG2 were labeled by red circle or pink dots to represent the structural differences. (D) The ADP binding sites in ARG2 from BTx623 and SC328C are different. The amino acids required for ADP binding were predicted based on the 3D structure of 3pfiB using COFACTOR and COACH by I-TASSER program and labelled as dots. The ARG2 tertiary structure in BTx623 was matched to the disease resistance RPP13-like protein 4 (ZAR1; template c6j5tC)

**Supplemental Figure S7. Phylogenetic analyses of ARG2 to related proteins in monocot plant species.**

Phylogenetic relationship between ARG2 and (A) 36related proteins from sorghum and other monocot species, (B) 17 closely related homologs from other monocot species. Phylogenetic trees constructed by MEGA X using the Maximum Likelihood method based on the JTT model with 1000 bootstrap replicates of amino acid sequences of ARG2 and selected species. Bootstrap percentages > 75% are shown at the nodes. *Sb*, *Sorghum bicolor*; *Sv*, *Setaria viridis*; *Ttu*,*Triticum turgidum; Ome*, *Oryza meyeriana*; *Osi*, Oryza sativa Indica Group; *At*, *Aegilops tauschii*; *Tti*,*Triticum timopheevii*; *Osj*,Oryza sativa Japonica Group; *Oo*, *Oryza officinalis*; *Op*,*Oryza punctat*e; *Ec*, *Eragrostis curvula*; *Si*, *Setaria italica*; *Hv*, *Hordeum vulgare*; *Omi*, *Oryza minuta*, *Bd*; *Brachypodium distachyon*; *Ob*,*Oryza brachyantha*; *Tu*,*Triticum urartu*; *Ath*, *Arabidopsis thaliana*. *Arabidopsis, dicot,* is included as an outlier control.

**Supplemental Figure S8. Phylogenetic relationship between ARG2 and functionally characterized resistant proteins from different plant species.**

Intact NB-ARC domains of the ARG2 and functionally characterized NLR genes from different plant species were used in the construction. Phylogenetic tree constructed by MEGA X using the Maximum Likelihood method based on the JTT model with 500 bootstrap replicates. Bootstrap percentages > 75% are shown at the nodes.

**Supplemental Figure S9. Metabolite analysis result of ARG2-mediated resistance, and analysis of growth and physiological traits among ARG2 NILs in the absence of the pathogen in a controlled environment phenotyping facility.**

(A) MetPA “MS peak to pathway” analysis of metabolites in *ARG2* NILs in response to Csgl1 at 48 hours post inoculation (hpi). Top hit metabolic pathways based on “mummichoh pathway enrichment” algorithm and gene set enrichment analysis (GSEA) using the KEGG *Oryza* metabolite database. Zeatin biosynthesis, Porphyrin and Chlorophyll metabolism, Tryptophan metabolism, Anthocyanin biosynthesis, Arginine and proline metabolism, Glucosinolate biosynthesis, Lysine degradation. (B) NILs in the growth chamber without pathogen inoculation (left) and on the navigation belt for imaging (right). (C) Anthocyanin reflectance, and (D) plant senescence reflectance indices at flowering time (97 days after planting, dap) and grain filling stage (119 dap), based on hyperspectral image. Estimates of (E) Top-average-hue throughout the growth period. rr, homozygote recessive; H, heterozygote; RR, homozygote dominant genotypes. (F) Correlation between physiological and growth traits estimates between the flowering time and the grain filling stage. SACH, Side-average-convex-hull; TPS, Top-plant-surface, TLA, Total-leaf-area; PRI, Phytochemical-reflectance-index; NRI, Nitrogen-reflectance- index; PSRI; ARI, Anthocyanin reflectance index; Plant-senescence-reflectance-index. The suffices “1” in the x-axis and “2” in the y-axis labels show the flowering time (97^th^ days after planting) and the grain filling stage (119 days after planting), respectively. ANOVA and Tukey HSD functions were used to test statistical significance of differences among genotypes.

**Supplemental Figure S10. Panicle exertion and leaf color under greenhouse growth conditions in the parental lines, and the near-isogenic lines that differ in ARG2 locus.**

(A) Leaf at maturity (B) Panicle exertion.

Supplemental File S1. Alignment of the *ARG2* nucleotide sequences from 15 sorghum lines.

The *ARG2* coding sequences in four resistant (SC328C, Keller, SC237 and Rio) and 11 susceptible lines were aligned. The alignment shows four different ARG2 alleles. *ARG2* allele in SC328C is similar to that in Keller, SC237 and Rio. The susceptibility ARG2 allele from SC265 is a unique allele. TAM428, BTx642, SC23, IS9830, PQ434 and IS18760 show the same allele. ICSV745, IS8525, Malisor84-7 carry the susceptibility allele like BTx623. The premature stop codons are shaded in red. The premature stop codons are shaded in red. These amino acid sequences were aligned using “multiple sequence alignment” tools: kalign [www.ebi.ac.uk/Tools/msa/kalign/] and mview [www.ebi.ac.uk /Tools/msa/mview/].

Supplemental File S2. Alignment of the *ARG2* amino acid sequences in 15 sorghum lines.

The ARG2 amino acid sequences in four resistant (SC328C, Keller, SC237 and Rio) and 11 susceptible lines were aligned. The alignment shows four different ARG2 alleles. *ARG2* allele in SC328C is similar to that in Keller, SC237 and Rio. The susceptible ARG2 allele from SC265 is a unique allele. TAM428, BTx642, SC23, IS9830, PQ434 and IS18760 show the same allele. ICSV745, IS8525, Malisor84-7 carry the susceptibile allele like BTx623. The premature stop codons are shaded in red. These deduced amino acid sequences were aligned using “multiple sequence alignment” tools: kalign [www.ebi.ac.uk/Tools/msa/kalign/] and mview [www.ebi.ac.uk /Tools/msa/mview/].

Supplemental File S3. Comparison of the ARG2 allele in SC265 and TAM428 (15.5% non-synonymous codon).

Supplemental File S4. Comparison of the amino acid sequences of the LRR domain of ARG2 in SC328C and BTx623.

**Supplemental Table S1**. Response of sorghum lines to five strains of *Colletotrichum sublineola*.

**Supplemental Table S2**. BSA-Seq analysis results.

BSA-seq results that show the SNP index from the resistant and the susceptible bulks of the ARG2 mapping population.

**Supplemental Table S3**. Sorghum variants that were evaluated for host response to Csgl1.

Genotyping by PCR was carried out using a size-marker in the *ARG2* mapping interval.

**Supplemental Table S4**. Conserved NLR amino acids that vary between the SC328C and BTx623 ARG2.

**Supplemental Table S5**. Additional metabolite analysis results.

All pathway hits in the Oryza KEGG based on “MS peak to pathway” function in MetPA analytical pipeline. All pathway hits with their corresponding matched compound features. Only the proton was chosen in “Edit currency metabolites” function due to the ionization mode that was used to generate the data. All other MetPA statistical parameters were under the default mode.

**Supplemental Table S6**. List of primers used in this study.

